# Restoring fertility in yeast hybrids: breeding and quantitative genetics of beneficial traits

**DOI:** 10.1101/2021.01.21.427634

**Authors:** S. Naseeb, F. Visinoni, Y. Hu, A. J. Hinks Roberts, A. Maslowska, T. Walsh, K. A. Smart, E. J. Louis, D. Delneri

## Abstract

Hybrids species can harbour a combination of beneficial traits from each parent and may exhibit hybrid vigour, more readily adapting to new harsher environments. Inter-species hybrids are also sterile and therefore an evolutionary dead-end unless fertility is restored, usually via auto-polyploidisation events. In the *Saccharomyces* genus, hybrids are readily found in nature and in industrial settings, where they have adapted to severe fermentative conditions. Due to their hybrid sterility, the development of new commercial yeast strains has so far been primarily conducted via selection methods rather than breeding. In this study, we overcame infertility by creating tetraploid intermediates of *Saccharomyces* inter-species hybrids, to allow continuous multigenerational breeding. We incorporated nuclear and mitochondrial genetic diversity within each parental species, allowing for quantitative genetic analysis of traits exhibited by the hybrids, and for nuclear-mitochondrial interactions to be assessed. Using pooled F12 generation segregants of different hybrids with extreme phenotype distributions, we identified QTLs for tolerance to high and low temperatures, high sugar concentration, high ethanol concentration, and acetic acid levels. We identified QTLs that are species specific, that are shared between species, as well as hybrid specific, where the variants do not exhibit phenotypic differences in the original parental species. Moreover, we could distinguish between mitochondria-type dependent and independent traits. This study tackles the complexity of the genetic interactions and traits in hybrid species, bringing hybrids into the realm of full genetic analysis of diploid species, and paves the road for the biotechnological exploitation of yeast biodiversity.

## Introduction

Hybridization is important evolutionarily, as well as industrially, as it may offer advantageous gene combinations and traits of interest to the newly formed hybrids. Hybrids are commonly found in both natural and domestic situations with as many as 25% of plant species and 10% of animal species hybridizing naturally (Mallet 2007). The genus *Saccharomyces* encompasses 8 species, included the newly discovered *S. jurei* (Naseeb et al, 2017 and Naseeb et al 2018), which all readily hybridize among them. In the *Saccharomyces* yeasts there are many examples of hybrids from both natural (wild) as well as fermentation sources, and indeed as many as 10% of all yeast isolates are hybrids (Liti et al. 2005; Liti and Louis 2005; Liti et al. 2006). The stringent condition of beer and wine fermentations in particular, represent a fertile ground for hybrids, influencing their creation, stabilisation, and phenotypic and genetic makeup (Bradbury et al. 2006; Gonzalez et al. 2006; Gonzalez et al. 2008; Lopandic 2018; Masneuf et al. 1998). Indeed, *S. pastorianus* (synonym *S. carlsbergensis*), a *S. cerevisiae/S. eubayanus* hybrid, has been employed in beermaking since the 15th century (Dunn and Sherlock 2008; Libkind et al. 2011; Nakao et al. 2009). *S. cerevisiae/S. kudriavzevii hybrids* have also been isolated from wine and cider fermentations along with *S. cerevisiae*/*S. uvarum* hybrids, and the triple hybrid between *S. cerevisiae*, *S. uvarum* and *S. kudriavzevii* (Gonzalez et al. 2006; Groth et al. 1999; Naumova et al. 2005). Moreover, examples of hybrids between the closely related species *S. cerevisiae* and *S. paradoxus* have been isolated from wild environments (Barbosa et al. 2016). Interspecies hybrids of *Saccharomyces* species have therefore been used as model organisms for studying natural evolution, speciation and fitness adaptation to different environments.

Recently, much work has gone into the generation of *de novo* yeast hybrids, exploiting their potential for the production of biofuels (Peris et al. 2017; Snoek et al. 2015), brewing (Krogerus et al. 2015; Mertens et al. 2015), and winemaking (Bellon et al. 2013; Garcia-Rios et al. 2018). Inter-species hybrids are not only selected for their capability to combine the advantageous traits of the parent strains, as the genomes from both parents undergo chromosomal rearrangements, mutations, wide spread transcriptional changes (Hewitt et al. 2020; Lairón-Peris et al. 2020), and gene loss and gene duplications which also impact the nature of protein complexes formed (Piatkowska et al., 2013). Hence, new and improved phenotypes can arise thanks to heterosis or hybrid vigour (Bradbury et al. 2006; Lopandic et al. 2016; Pfliegler et al. 2012; Sipiczki 2008).

A great deal has been learned on the acquired properties of particular hybrids from comparative genomics and molecular studies. However, to date, no thorough genome-wide analysis of genetic contributions to traits of industrial interest has been possible. The sterility of the inter-species yeast hybrids is in fact hindering the application of both predictive quantitative approaches and any attempt of strain improvement via breeding. A previous study by Greig et al. (2002) showed that sterility of inter-species yeast hybrids can be overcome by creating tetraploids, using a MAT-locus deletion strategy. The engineered tetraploid hybrids were able to produce viable diploid progeny after undergoing meiosis (Greig et al. 2002). Another study by Schwartz et al (2012) demonstrated the ability to mate a hybrid for quantitative trait loci (QTL) analysis using expression of *HO* to switch a diploid non-mater to a mater. QTL mapping in particular has shown to be a powerful tool for understanding the genetic basis of various complex traits and has been similarly applied in industrial and medical applications (Deutschbauer and Davis 2005; Steinmetz et al. 2002; Zimmer et al. 2014). Nonetheless, QTL analysis and understanding of hybrid genetics remains poor and limited to a single generation of meiosis in these studies, due to the sterility of *Saccharomyces* hybrids.

In this study, we used two different approaches to bring *de novo* yeast hybrids into the realm of genetic analysis and improvement by breeding. A large set of *de novo* hybrids were engineered by crossing geographically distinct *Saccharomyces* strains of different species. We were able to generate genetically and phenotypically diverse populations through multiple rounds of interbreeding. Diploid F2 and F12 progeny were phenotyped for several traits, and two approaches towards genotyping pools of phenotypic extremes were used to map genetic variation in both parental genomes responsible for the phenotypes selected. For 3 sets of hybrids, F12 diploid hybrid progeny were arrayed and phenotyped under several different conditions. For each hybrid, two versions were assessed, one each with each parental species mitochondrion. The top 20 and bottom 20 from an array of 384 were pooled for sequencing and the frequencies of segregating SNPs in the two parental species genomes were compared to identify QTL involved in the traits. For one of these hybrids (2 mitochondrial versions), strong selection was applied to a population of 10^8^ F12 progeny resulting in fewer than 10^6^ survivors whose pooled genomic DNA was compared to the pool prior to selection, for several conditions. We demonstrate *Saccharomyces* hybrids are now amenable to all the tools available for yeast, including breeding and utilisation of the vast genetic diversity available. We show that there are QTL both unique to one parent species or shared by both, QTL dependent and independent of the mitochondrial origin, and QTL that are specific to the hybrid and not to the parent species where the variants are segregating.

## Results and Discussion

### Construction of tetraploid yeast hybrids from a variety of parental strains leads to restoration of fertility

Hybrid sterility can be overcome by doubling the genome of the hybrid. Such fertility restored hybrids, known as amphidiploids, are commonly found in plants and represent the majority of major evolutionary events in angiosperms (Moghe et al. 2014; Wang et al. 2011; Zhang et al. 2016). Tetraploidisation, resulting in amphidiploids, can also restore fertility in yeast hybrids (Greig et al. 2002).

We constructed yeast hybrid tetraploid lines combining four different genomes of strains coming from two separate species belonging to the *Saccharomyces* genus. To construct such fertile tetraploid hybrids, we employed two different strategies (Figure 1). First *inter-species* diploid hybrids were constructed (5 for *S. cerevisiae/S. jurei* and 10 for *S. cerevisiae/S. kudriavzevii*, Table S1), followed by reciprocal deletion of the MAT locus, and subsequent crossing of the two diploid mater hybrids. In the second strategy *intra-species* diploid strains were first constructed by crossing diverged populations of the same species (10 *S. cerevisiae*, 6 *S. uvarum* and 2 *S. eubayanus* intra-species crossings, Table S2), followed by reciprocal deletion of the MAT locus, and subsequent hybridization of the two diploid maters. Tetraploid lines of *S. cerevisiae*/*S. kudriavzevii* (Sc/Sk) and *S. cerevisiae*/*S. jurei* (Sc/Sj) hybrids were created with this first strategy while *S. cerevisiae*/*S. uvarum*, and *S. cerevisiae*/*S. eubayanus* were constructed using the second approach.

**Figure 1.**
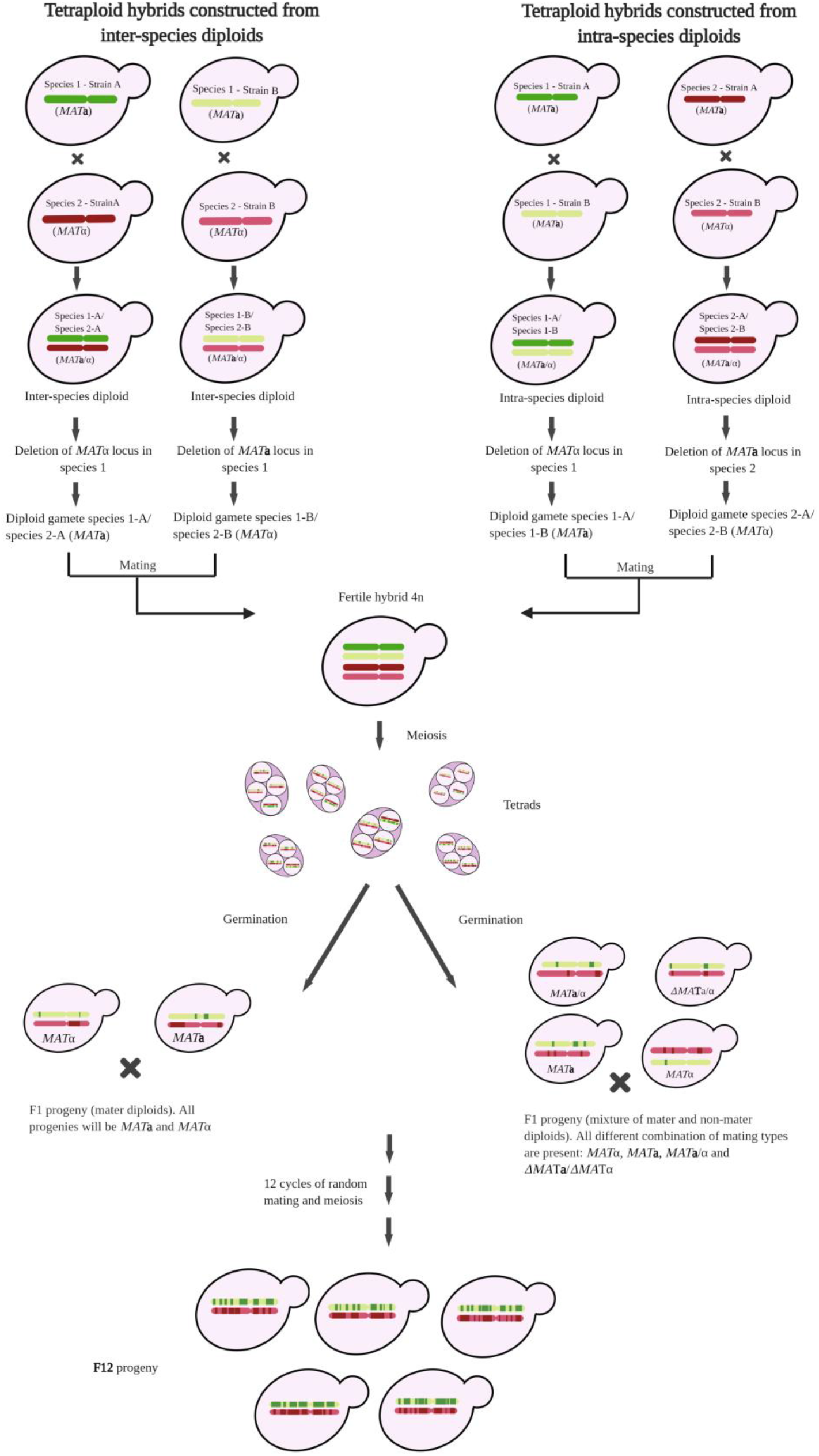
Construction of fertile hybrids. The tetraploid hybrids were constructed using two different strategies. In the first strategy, two different *Saccharomyces sensu stricto* species were crossed to obtain 2n *inter-species* diploid hybrids (species 1 and species 2 are represented in shade of green and red, respectively). Two different of such hybrids were made to behave as gametes by deleting either the *MAT***a** or α locus in one species and subsequently crossed to make the fertile tetraploid hybrid (4n). In the second strategy, two diverged populations (A and B) of the same species were crossed to construct the 2n *intra-species* hybrids. Two different intra-species lines were also made to behave as gametes by deleting the *MAT***a** from species 1 and *MAT*α from species 2, and subsequently these were crossed to construct the 4n hybrids. The tetraploid hybrids were sporulated and germinated to obtain the diploid F1 progeny which were randomly mated and sporulated several times until a F12 generation with a high level of scrambled genomes.

Hybrids between *Saccharomyces* species are homoplasmic and tend to carry mitochondrial DNA from only one parent. The natural hybrid *S. pastorianus* (hybridized from *S. cerevisiae* and *S. eubayanus*) carries the *S. eubayanus* mitochondria (E. Baker et al. 2015; Okuno et al. 2016), while other industrial hybrids of *Saccharomyces* species used in wine and cider production have retained *S. cerevisiae* mitochondrial genome (Masneuf et al. 1998). It has also recently been shown that the type of mitochondria inherited affects the phenotype (Albertin et al. 2013; E. P. Baker et al. 2019; X. C. Li et al. 2019) and the transcriptional network (Hewitt et al. 2020) in hybrids. Therefore, here, each of the tetraploid hybrids constructed were also selected for different mitotypes. Throughout the paper, in the hybrid nomenclature, a subscripted _m_ following the initial of the species, represents the particular mitochondria inheritance of that hybrid (*i.e.* Sc_m_/Sj is tetraploid hybrid containing Sc mitochondria, Sc/Sj_m_ is tetraploid hybrid containing Sj mitochondria).

A total of 226 tetraploid hybrids were created and each hybrid had four unique parental strains and a unique mitochondrion contributing to the genome (Table S3). All the constructed tetraploids were fertile as they had a homologous set of chromosomes to align in meiosis. While diploid hybrids between species of *Saccharomyces* genus are reproductively isolated (Liti et al. 2006; Naseeb et al. 2017), the tetraploid hybrids constructed here, were fertile and exhibited spore viability as high as 98% (Table S4). Ultimately, the diploid F1 spores obtained from the tetraploid hybrids were sequentially backcrossed and sporulated eleven times until the F12 generation (Figure 1). From the F12 generation around 384 spores for each were isolated for further studies.

### Meiotic offspring of tetraploid hybrids exhibit broad phenotypic diversity

Inter-crossing different populations of *Saccharomyces sensu stricto* yeast species over many generations reduces linkage disequilibrium by increasing recombination. To assess whether the fitness traits are associated with genetic linkage, we assessed the phenotypic landscape of F1 and F12 diploid generations in up to 12 conditions encompassing different growth temperatures, carbon sources and stressors. An example of phenotypic divergence between F1 diploid segregants of tetraploid Sc/Sj (OS3/D5088/OS104/D5095) and Sc/Sk (OS253/OS575/OS104/IFO1802) harbouring different mitotypes is reported in Figure S1. Significant fitness differences were seen in all the segregant lines with a dispersion up to 0.33 (quartile coefficient of dispersion, Table S5), with some progeny being fitter than any of the parents (Figure S2) and some being less fit (transgressive variation) in virtually all cases.

When colony size is normalised within each specific condition, to allow the teasing apart of fitness differences between spores derived from the same tetraploid line, F12 segregants of Sc_m_/Sj, Sc/Sj_m_, Sc_m_/Sk, and Sc/Sk_m_ again exhibited a large phenotypic variation in all the tested conditions (Figure 2).

**Figure 2.**
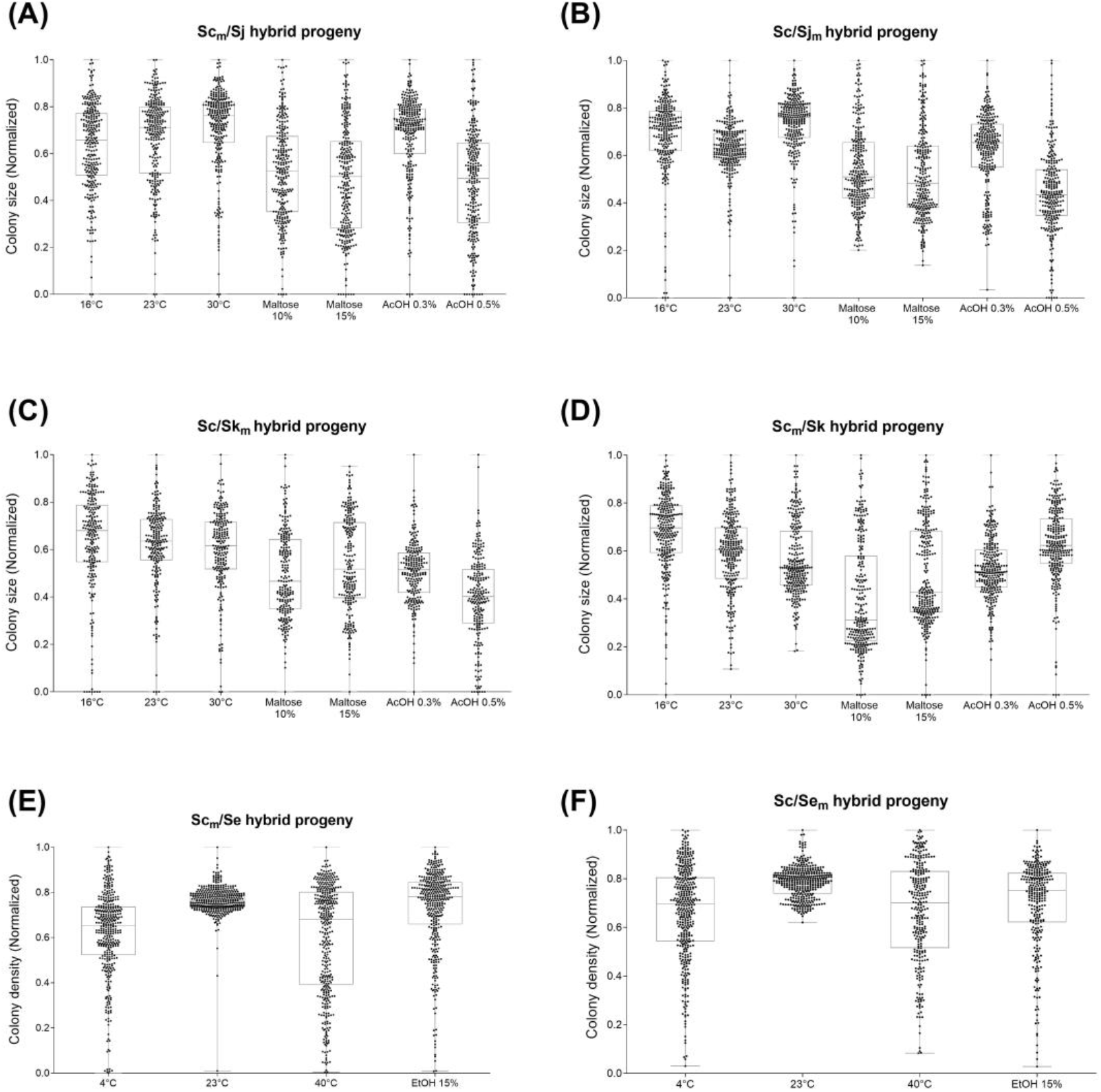
A box plot of the fitness of F12 diploid progeny for *S. cerevisiae/S. jurei* (Scm/Sj and Sc/Sjm) and *S. cerevisiae/S. kudriavzevii* (Scm/Sk and Sc/Skm) and *S. cerevisiae/S. eubayanus* (Scm/Se and Sc/Sem) hybrids. For Sc_m_/Sj (panel A), Sc/Sj_m_ (panel B), Sc_m_/Sk (Panel C) and Sc/Sk_m_ (Panel D), the normalised colony size was used as proxy of fitness (see Material and Methods) and was scored in YPD at different temperatures 16°C, 23°C, and 30°C, in YP-Maltose (10% and 15%), and in YPD with 0.3% and 0.5% acetic acid (AcOH). For Sc_m_/Se (Panel E) and Sc/Se_m_ (Panel F) the colony optical density was used as proxy of fitness (see Material and Methods) and was scored in YPD at different temperature 4°C, 23°C, and 40°C, and in YPD with 15% Ethanol (EtOH). Each black dot represents a different F12 diploid hybrid. The upper and lower error bars represent the minimum and maximum values.

Given that the tetraploid lines are composed of *S. cerevisiae* combined with genomes of other *Saccharomyces* species with different levels of phylogenetic distance, we investigated whether such differences in genome divergence has an impact on phenotypic plasticity in any given condition. By analysing the un-normalised fitness data, to tease apart differences in the colony size range between progeny from separate tetraploid lines, no striking differences were found in either range or dispersion between different hybrids (Figure S3). Therefore, the different levels of divergence of the genomes present in the hybrid lines does not impact significantly on phenotypic range and plasticity in the progeny.

Twenty F12 individuals with high fitness, and 20 with low fitness at low temperature, high maltose and high acetic acid conditions, were then chosen to carry out QTL analysis. Such conditions are relevant to fermentation industries: low temperature is required for the storage of brewing yeast and also for fermentation of lager beer; maltose is one of the key wort sugars and the high concentration of this sugar mimics the osmotic pressure exerted upon yeast in high gravity wort (Casey et al. 1984; Gibson et al. 2007); acetic acid is found in grape must and wine producing yeasts, are known to require the resistance to this stress (Belloch et al. 2008; Gibson et al. 2007).

### Phenotypic diversity of tetraploid hybrids is underpinned by the presence of QTL

To identify the genetic basis underlying the observed phenotypic diversity we performed QTL analysis on selected segregant pools of Sc/Sj, Sc/Sk, Sc/Se and Sc/Su hybrids. Two different methods for QTL analysis were employed. The Multipool technique, pioneered by (Edwards and Gifford 2012), was used to analyse the F12 generation of Sc/Sj, Sc/Sk, and in Sc/Se hybrids, while the Pooled Selection method, or bulk segregant analysis (Ehrenreich et al. 2010; Parts et al. 2011) was applied to Sc/Se and Sc_m_/Su hybrids.

Both approaches proved successful in mapping QTL regions in a variety of conditions for all hybrids, however a higher number of QTLs and more consistent results across the conditions tested were obtained using the Multipool approach (Table S6). Our results indicate that the Multipool approach with individuals at each extreme of a phenotype distribution is more efficient than a highly selected pool approach. However, the highly selected pool approach can identify rare genotypes of linked recombinant variants that are too rare to be in the arrays used to choose individuals for the Multipool approach.

From segregants generated from the tetraploids Sc_m_/Sj and Sc/Sj_m_, a total of 56 QTLs were identified in the *S. cerevisiae* genomes with an average length of 19.4 kb (Table 2). Despite the high similarity between the two *S. jurei* parental strains (Naseeb et al. 2018), we were able to map 62 QTL regions in the genome of this species. However, with an average length of 35.5 kb, the QTL mapping intervals were the longest observed in the hybrids generated due to the lower density of segregating markers in the two genomes of this species.

An even higher number of QTL was mapped from the progeny of Sc_m_/Sk and Sc/Sk_m_ tetraploids with up to 155 and 128 regions mapped in *S. cerevisiae* and *S. kudriavzevii*, respectively (Table 2). Here, the QTL intervals were narrower than those seen in the *S. jurei* genome, with only 15.6 kb average length on *S. cerevisiae* alleles and 18.6 kb on *S. kudriavzevii* ones as there is a higher density of segregating markers in these genomes.

Similar results were obtained in Sc_m_/Se and Sc/Se_m_ hybrids analysed for their fitness in high ethanol concentration and at high and low temperatures (40°C and 4°C) (Table 2). Here, we were able to identify 111 and 64 QTLs regions in *S. cerevisiae* and *S. eubayanus*, respectively, with an average length of 18.5 kb and 21.82 kb.

A total of 28 genes mapped in different QTL in Sc/Sj and Sc/Sk hybrids were classified as potential causal genes, as their role in the selection condition was already confirmed by previous published work (Cherry et al. 2012) (Table S7). For example, one of the acetic acid QTL detected in *S. cerevisiae* chromosome III (52-97 kb) in Sc_m_Sj hybrids contains variant alleles of *LEU2*. The gene encodes for a β-isopropyl-malate dehydrogenase and null mutants are reported as sensitive to acetic acid while its overexpression increase acetic acid resistance (Hueso et al. 2012).

Amongst the genes identified, a total of 43 genes with segregating alleles found in low temperature QTL were previously identified in large-scale competition studies carried out in *S. cerevisiae* at 16°C, with an additional 5 being described as cold favouring by thermodynamic model predictions (Table S7) (Paget et al. 2014).

Thanks to the abundance of data on heat and ethanol sensitivity in *S. cerevisiae* (Auesukaree et al. 2009; Cubillos et al. 2013; Parts et al. 2011; Sinha et al. 2008; Teixeira et al. 2009), a high number of potential casual genes with segregating variation were identified in the QTL regions of Sc/Se hybrids. Thus, we were able to identify up to 38 genes in the 44 QTL regions for the *Sc* alleles that likely promote a fitness advantage while growing at 40°C (Table S7). Amongst these, *IRA1*, a regulator of the RAS pathway, was previously validated in previous studies as a heat-QTL in OS3/OS104 crosses (Parts et al. 2011). Moreover, two additional genes involved in the RAS/cAMP signalling pathway (*ESB1* and *GPB2*) were mapped in heat QTLs, supporting its involvement in in mediating heat resistance as previously suggested by Parts et al. (2011).

The potential causal genes detected in *S. cerevisiae* genomes in Sc/Sj, Sc/Sk, and Sc/Se hybrids may contain amino-acid variants that are affecting protein function. Hence, we analysed these genes through SIFT analysis (Sorting Intolerant From Tolerant), to identify non-synonimous SNPs underlying the observed phenotypic difference between alleles (Kumar et al. 2009, Bergstrom et al. 2014). SIFT analysis was carried out on the 79 predicted *S. cerevisae* casual genes. A strong effect on the protein function was detected in 24% of potential causal genes due to amino acid differences between the *S. cerevisiae* parental strains (Table S8). The predicted effect of the SNPs in these allelic variants further supports the efficacy of our approach.

GO analysis did not help to narrow down choices of potential casual gene candidates, since the enrichment GO terms were at broad level only generally associated with intracellular membrane bound organelle, cytoplasm, catalytic activity, and cellular processes in all the conditions.

In total 14, 22 and 11 pleiotropic QTLs were mapped in Sc/Sj, Sc/Sk and Sc/Se hybrids, respectively (Table S9). A 7 kb region on the *S. cerevisiae* chromosome XIII was common across all conditions tested for ScSj_m_ hybrids, but, interestingly, it was not detected for Sc_m_Sj in any condition tested. This region contains the genes *CLU1* (a subunit of eIF3), *ANY1* (a protein involved in phospholipid flippase) and *HXT2* (a high-affinity glucose transporter). It is possible that the phenotypic effect of variation in these genes depends solely on mitochondrial-nuclear interactions, independent of the condition. *CLU1* is known to play a role in mitochondrial distribution and morphology but it maintains its respiratory function and inheritance (Dimmer et al. 2002; Fields et al. 1998). The Δ*clu1* mutants possess a more condensed mitochondrial mass found at one side of the cell (Fields et al. 1998). They are haplo-insufficient in nutrient limited media (Fields et al. 1998; Pir et al. 2012) and haplo-proficient in phosphorus limited media (Delneri et al. 2008).

In parallel to the Multipool, pooled selection experiments were performed on Sc_m_/Se and Sc/Se_m_, Sc_m_/Su segregants. Specifically, Sc_m_/Se and Sc/Se_m_ were selected for a variety of selectable traits of industrial relevance, spanning from high maltose or glucose concentrations (35%), to H_2_O_2_ treatment (4 mM), and very low temperature (4°C); while the Sc_m_Su hybrid progeny were selected in YPD at high temperature (40°C) and YPD with levulinic acid (50 mM), acetic acid (0.35%) at 23°C.

The linkage analysis on the pooled selection segregants yielded narrow intervals, averaging at 18.1 kb and 20 kb in Sc/Se and Sc_m_/Su hybrids, respectively. However, only a limited number of QTL were identified, except for YP-Maltose (Table S6). No QTL were found among segregating *S. uvarum* alleles (Table S10).

35 of the 41 QTL regions detected in Sc_m_/Se and Sc/Se_m_ segregants were present in hybrids with a fitness advantage in high maltose concentrations (Table S10). However, it was unexpected that only one QTL was identified when selection took place at 4°C, given that in the multipool analysis in the same condition, 45 QTLs were identified. Characteristics such as growth at low temperature, which, albeit selectable, do not show extremes phenotypes, are better discriminated using the Multipool approach as it exploits the richness of fitness data acquired through individual phenotyping. Growth at 4°C may not be strong enough selection, *i.e.* doesn’t kill the majority of individuals in the population, resulting in less discrimination in the pooled selection approach.

Similarly, pooled selection of Sc_m_/Su segregants yielded only 9 intervals of interest, all mapping to *S. cerevisiae* alleles across the four conditions tested. These regions included a single interval conferring tolerance to acetic acid, four conferring selective advantage at high temperature, and four giving tolerance in an environment containing levulinic acid. As mentioned above none were found in the *S. uvarum* genome.

As with Multipool QTLs, we identified several genes, which had already been reported as having a phenotypic effect or a function closely linked to the condition tested. For Sc_m_/Se and Sc/Se_m_ hybrids, we detected in the Pooled Segregants in high maltose concentrations 8 genes where deletion was reported to cause osmotic stress sensitivity (Table S11). Among these, *SKO1*, a basic leucine zipper transcription factor, mapped in *S. eubayanus* alleles of Sc/Se_m_ segregants, has been described having a major role in mediating HOG pathway-dependent osmotic regulation (Rep et al. 2001).

### Different types of mitochondria have a profound effect on the QTL landscape

Mitochondrial-nuclear interactions have been reported as having a major role in phenotypic variation both in intra-species and inter-species yeast hybrids (E. P. Baker et al. 2019; Chou et al. 2010; Hewitt et al. 2020; Hsu and Chou 2017; Lee et al. 2008; X. C. Li et al. 2019; Paliwal et al. 2014), affecting respiration (Albertin et al. 2013) fermentation properties (Hewitt et al. 2020; Solieri et al. 2008), progeny fitness (Zeyl et al. 2005; Zeyl 2006), reproductive isolation (Chou and Leu 2010; Lee et al. 2008; Szabo et al. 2020) and nuclear transcription (Hewitt et al. 2020). Studies on inter-species yeast hybrids in the lab have shown a correlation between the origin of the mitochondrial genome and the higher stability of one of the nuclear genomes (Antunovics et al. 2005; Karanyicz et al. 2017; Peris et al. 2020). Moreover several mitochondrial-nuclear incompatibilities leading to respiratory deficiencies have been identified in yeast hybrids (Chou et al. 2010; Lee et al. 2008). The incompatibility of the *S. uvarum* mitochondrion with the *S. cerevisiae* nucleus was reinforced by the transplacement of mitochondria isolated from *S. uvarum* (Osusky et al. 1997; Spirek et al. 2000). Mitochondrial-nuclear incompatibilities are also associated with the splicing of mitochondrial intron cox1I3β (Spirek et al. 2014).

Mitochondrial-nuclear epistasis has been shown to affect phenotypes in several taxa. In insects, such as *Drosophila* and *Callosobruchus*, exchange of mtDNA variant has led to decreased metabolic rate (Dowling et al. 2007) and shortened life span (Clancy 2008; Zhu et al. 2014). In mice it is known to affect cognition and respiratory functions (Betancourt et al. 2014; Roubertoux et al. 2003). Interactions between mitochondrial and nuclear genomes can result in cytoplasmic male sterility in plants (Hu et al. 2014) and impact ageing and longevity in humans (Tranah 2011; Wolff et al. 2014). In yeast, mitochondrial-nuclear epistasis contributes to phenotypic variation and coadaptation to changing environmental conditions (De Chiara et al. 2020; Paliwal et al. 2014; Wolters et al. 2018; Zeyl et al. 2005). However, the complexity and the diversity of the hybrid background poses a challenge to a thorough mapping of the epistasis in genome-wide studies. Here, in order to evaluate how the mitochondria inherited may affect the QTL landscape, we compared QTL regions mapped in diploid hybrid progeny derived from tetraploid lines harbouring different mitochondria but the same parental nuclear inputs.

Interestingly, the majority of QTL detected via the Multipool method were exclusive to a specific mitotype (Figure 3). In fact, in all of the conditions analysed, only ca. 2.45% QTL regions (the mean percentage for all hybrids in all the conditions) were in common among the segregants from tetraploid hybrids with different mitochondria. This difference in the QTL landscape between the same yeast hybrid cross, differing only for the mitotype, reinforces the idea that mitochondrial-nuclear interactions have a genome-wide effect and are important in the context of evolution, as already demonstrated by several studies both in yeast (Chou and Leu 2010; Lee et al. 2008; Vijayraghavan et al. 2019; Wolters et al. 2018) and in other organisms (Barrientos et al. 1998; Ellison and Burton 2006; Ellison et al. 2008; McManus et al. 2019; Mossman et al. 2016; Mossman et al. 2017; Wernick et al. 2019).

**Figure 3.**
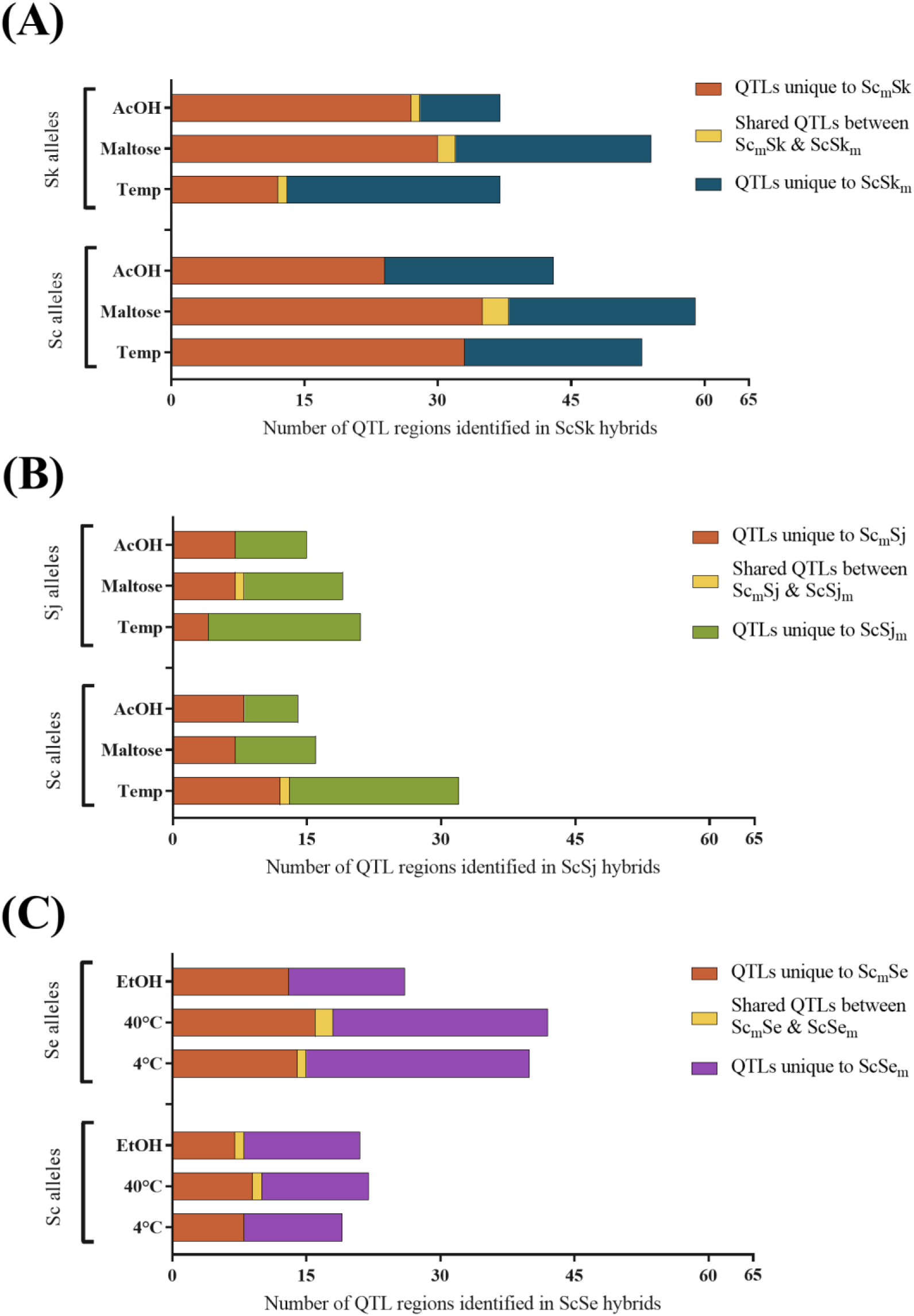
Hybrids with different mitotype exhibit a different QTL landscape. Boxplot of the distribution of QTLs in *S. cerevisiae/S. kudriavzevii* (Panel A), *S. cerevisiae/S. jurei* (Panel B) and *S. cerevisiae/S. eubayanus* hybrids (Panel C). The QTL regions are grouped first by the alleles and then by growth condition in which they were identified: acetic acid (AcOH), high maltose concentrations (Maltose), low temperature (16°C or 4°C), heat and ethanol (EtOH). The small proportion of QTLs shared between mitotypes of the same hybrid is represented in yellow.

The experiments carried out via Pooled Selection also demonstrated mitotype-specific QTLs. Only one QTL region was in common in the Sc/Se hybrids with different mitochondria: a 21 kb region on chromosome I, containing the genes *FLO1* and *PHO11* where specific Sc alleles are associated with an increase in fitness at high maltose concentrations. Given these are located in a subtelomeric region, known to be highly variable in copy number and location in addition to sequence, as well as difficult to assemble (Liti et al. 2009; Louis 2014), the causal genetic variation may be an unknown sequence linked to the *FLO1* and *PHO11* genes, as these regions are not assembled in all of the genomes utilized here.

Amongst the QTLs shared between Sc_m_Sj and ScSj_m_ segregants, a major maltose QTL near the telomeric regions of chromosome II of *S. cerevisiae* (780 – 795 kb) was detected with a high LOD score (> 17). The recurrence of this QTL and its high LOD score could be ascribed to the *MAL3* multigene complex in the sub-telomeric region and the natural variation in copy number, location and sequence of this complex (Liti et al. 2009). Furthermore, this maltose QTL is also mapped on the *S. cerevisiae* alleles of both Sc_m_/Sk and Sc/Sk_m_ hybrid pointing to a more general effect of these allelic variants rather than a strain background dependent one.

### Overlap of QTL regions between different hybrids facilitates the identification of causal genes

One way to help to identify relevant genes within QTL regions, at least for QTL not specific to a particular hybrid combination, is to investigate overlapping QTL regions detected in different hybrids. Such an approach can lead to the unambiguous identification of genes underlying the phenotypic effect observed.

Twelve overlapping intervals, shared by at least three hybrid genomes, were mapped in low temperature-, high maltose-, and high ethanol-QTLS, while an additional 52 intervals were overlapping between two hybrids (Table S12). Within acetic acid-QTLs, only 12 overlapping intervals were detected with no region shared between more than two species, suggesting a greater diversity of variants connected with the phenotype. Amongst these intervals, we identified several candidates with biological functions closely linked to the selection condition. For instance, a low temperature-QTL mapped both in *S. cerevisia*e and *S. kudriavzevii* includes *OSH6* (Figure 4 A), related to sterol metabolism and, as such, membrane fluidity, often considered a feature of low-temperature adaptation (Buzzini et al. 2012). Similarly, an acetic acid-QTL mapped in both Sc/Sj_m_ and Sc_m_Sk hybrids includes *NQM1*, a nuclear transaldolase involved in oxidative stress response, known to be induced by acetic acid stresses (Ludovico et al. 2001; Michel et al. 2015) (Figure 4 B).

**Figure 4.**
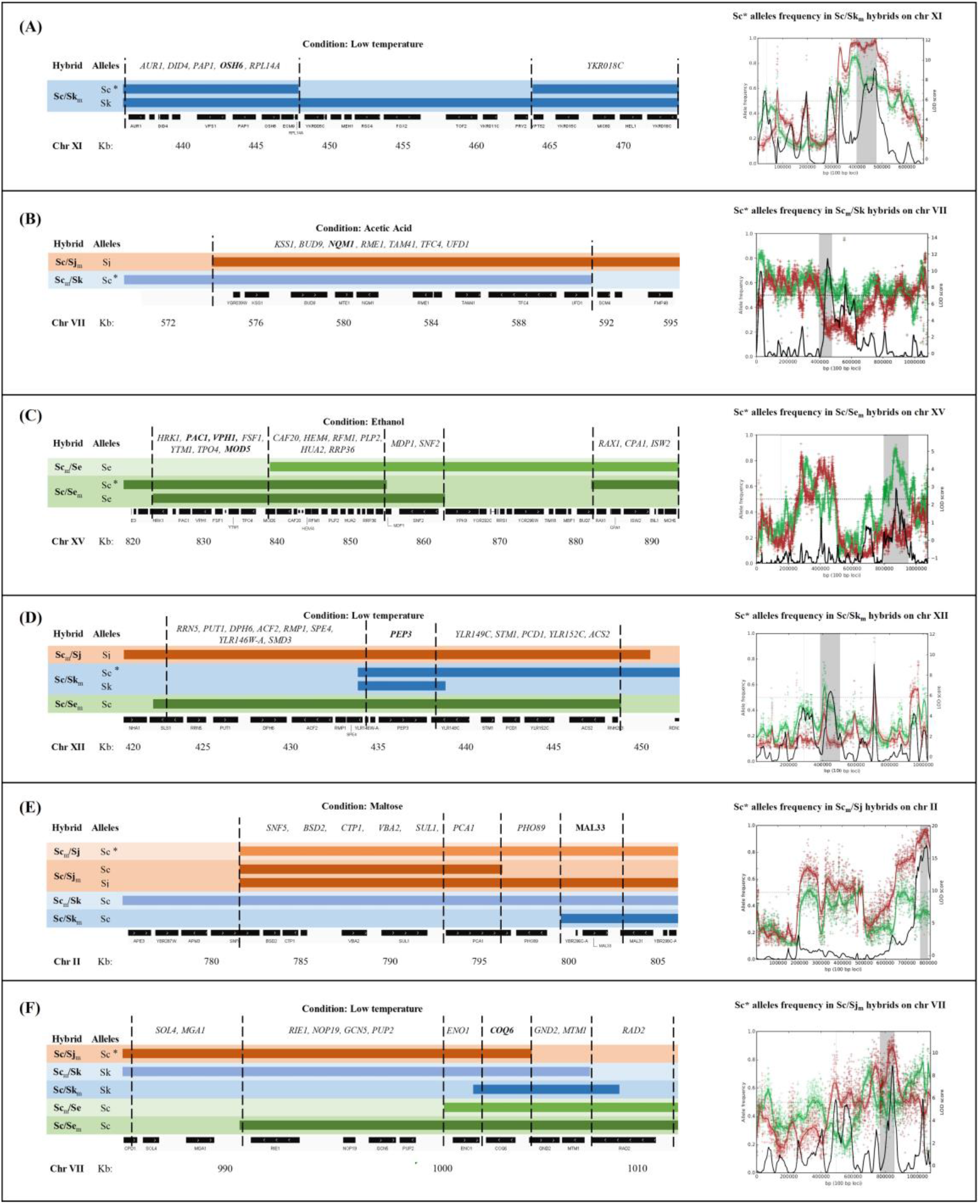
Example of inter-species QTLs detected in *S. cerevisiae/S. jurei*, *S. cerevisiae/S. kudriavzevii* and *S. cerevisiae/S. eubayanus* hybrids. The QTL regions, represented by coloured bars, are mapped onto *S. cerevisiae* chromosomes to identify the overlapping QTL intervals and the genes shared between different species and hybrids. The parental species alleles for each hybrid are stated and the genes in bold denotes potential casual genes. The QTL plots represents the frequency of *S. cerevisiae* alleles, marked in the figure with an asterisk, from different hybrid pool lines. The red and green lines represent the allele frequency of high and low fitness pools. Black line indicates the LOD score and grey area is the 90% credible interval of the significance. The dash line is the threshold of LOD considered in this study.

Ethanol- and high temperature-QTLs were analysed only in Sc/Se hybrids, limiting the outcomes compared to the other traits. Nonetheless, we found 3 overlapping ethanol-QTLs with a major region mapped on chromosome XV shared between genomes (Figure 4 C). Moreover, this region included the genes *PAC1*, *VPH1* and *MOD5* where null mutations are linked to a decreased fitness in ethanol supplemented media (Qian et al. 2012; van Voorst et al. 2006; Yoshikawa et al. 2009).

A major overlap was detected in the sub-telomeric region of chromosome II, with a maltose-QTL shared between the *S. cerevisiae* allele of all Sc/Sj and Sc/Sk hybrids and the *S. jurei* alleles of Sc/Sj_m_. The QTL contains a causal gene, *MAL33*, a MAL-activator protein, and two transporter *CTP1* and *PHO89*, involved in the transport of citrate and phosphate, respectively, which are key metabolites for the glycolytic pathway (Figure 4E).

Remarkably, two low temperature-QTLs regions were mapped in at least one mitotype of all tetraploid hybrids analysed through Multipool. The overlapping regions resulted in both cases in a single-gene intersection with *COQ6* (Figure 4F) and *PEP3* (Figure 4D) identified in 4 and 5 different intervals, respectively. *COQ6* is a mitochondrial monooxygenase which, in addition to its known role in mitochondrial respiration, is involved in fatty acid β-oxidation (Awad et al. 2018). *PEP3*, instead, is a component of the CORVET membrane tethering complex, and its role at cold temperature was previously suggested by large-scale competition studies (Paget et al. 2014). Moreover, SIFT analysis performed with mutfunc (Wagih et al. 2018) predicted a strong deleterious effect of the SNP in the OS253 *PEP3* variant, due to a substitution in a conserved region.

Overall, the overlaps of QTL regions among progeny from different tetraploid hybrids allows us to focus on the genes that are more likely to be responsible for the phenotypic variation. This comparison identifies alleles which exert a background-independent effect.

### Validation of QTL via reciprocal hemizygosis analysis

The narrow mapping intervals identified in the Sc/Sk_m_ hybrid allowed single-gene studies to validate the effect of candidate alleles, which were not strongly identified as casual genes. The fitness of the allelic variants was tested with reciprocal-hemizygosity analysis (RHA) (Steinmetz et al. 2002) performed on *ASI2*, *FUS3* and *GIT1*, which are candidate QTLs in acetic acid, low temperature (16°C) and high maltose, respectively (Figure 4, Panel A, Table S13).

Amongst the genes included in the acetic acid-QTLs, *ASI2*, part of the Asi ubiquitinase complex, was defined as a potential candidate gene as its null mutant was previously described as sensitive to oxidative stress in systematic studies (Higgins et al. 2002). *FUS3*, a MAPK protein, was previously described as haplo-insufficient in large-scale competition studies at 16°C (Paget et al. 2014). Lastly, we selected the plasma membrane permease *GIT1* included in a high LOD QTL region in chromosome III identified at high maltose concentrations, along *HMRA1*, *HMRA2*, *CDC39*, *CDC50* and *OCA4*. *GIT1* was deemed the most promising candidate as it is involved in phosphate and glycerol-3-phospate transport, important metabolites of the glycolytic pathway (Popova et al. 2010).

The phenotypic effect of *ASI2*, *FUS3* and *GIT1* alleles were validated via RHA performed on the hemizygote tetraploid parents (OS253/OS575/OS104/IFO1802) in YPD + 0.3% acetic acid at 30°C, YPD at 10°C and YP-maltose (15%) at 25°C, respectively (Figure 5, Panel B, C, D, Table S14). We observed a significant difference of the growth curve integral area (Table S14) between the performance of the allelic variants in all the three genes tested confirming their impact in the selection condition and validating them as having causal variant alleles. The *FUS3*^OS253^ and *GIT1*^OS104^ alleles performed better than the *FUS3*^OS104^ and *GIT1*^OS253^ alleles in terms of specific growth rate (p value = 0.0010 and 0.0335, respectively) and integral area (p value = 0.0047, 0.0435, respectively) (Table S14), mirroring what we have seen in our multipool QTL screening. For the *ASI2* gene, the *ASI2*^OS253^ was the allele prevalent in the high-fitness pool exposed to high concentration of acetic acid. In the phenotypic validation the *ASI2*^OS253^ variant performed worse than the *ASI2*^OS104^ in terms of integral area (p value = 0.0479) and T_mid_ (p value = 0.0182) but it reached a higher maximum biomass (p value = 0.0001). Although the growth parameters for these two alleles are clearly different, it is more ambiguous whether their fitness performance overlaps with that one detected in the QTL study where the *ASI2*^OS253^ was the allele prevalent in the high-fitness pool. This discrepancy could be due to other genes in the close proximity masking its effect or to the difference in the phenotypic screening employed, as the F12 segregants were assayed for their growth in solid media. Many of yeast QTLs have shown to be environment and background dependent and have linked sets of quantitative trait nucleotides (QTN) (Cubillos et al. 2011; Liti and Louis 2012; Sinha et al. 2006). In fact, it has been shown that sporulation efficiency in yeast is controlled by four QTNs (Gerke et al. 2009).

**Figure 5.**
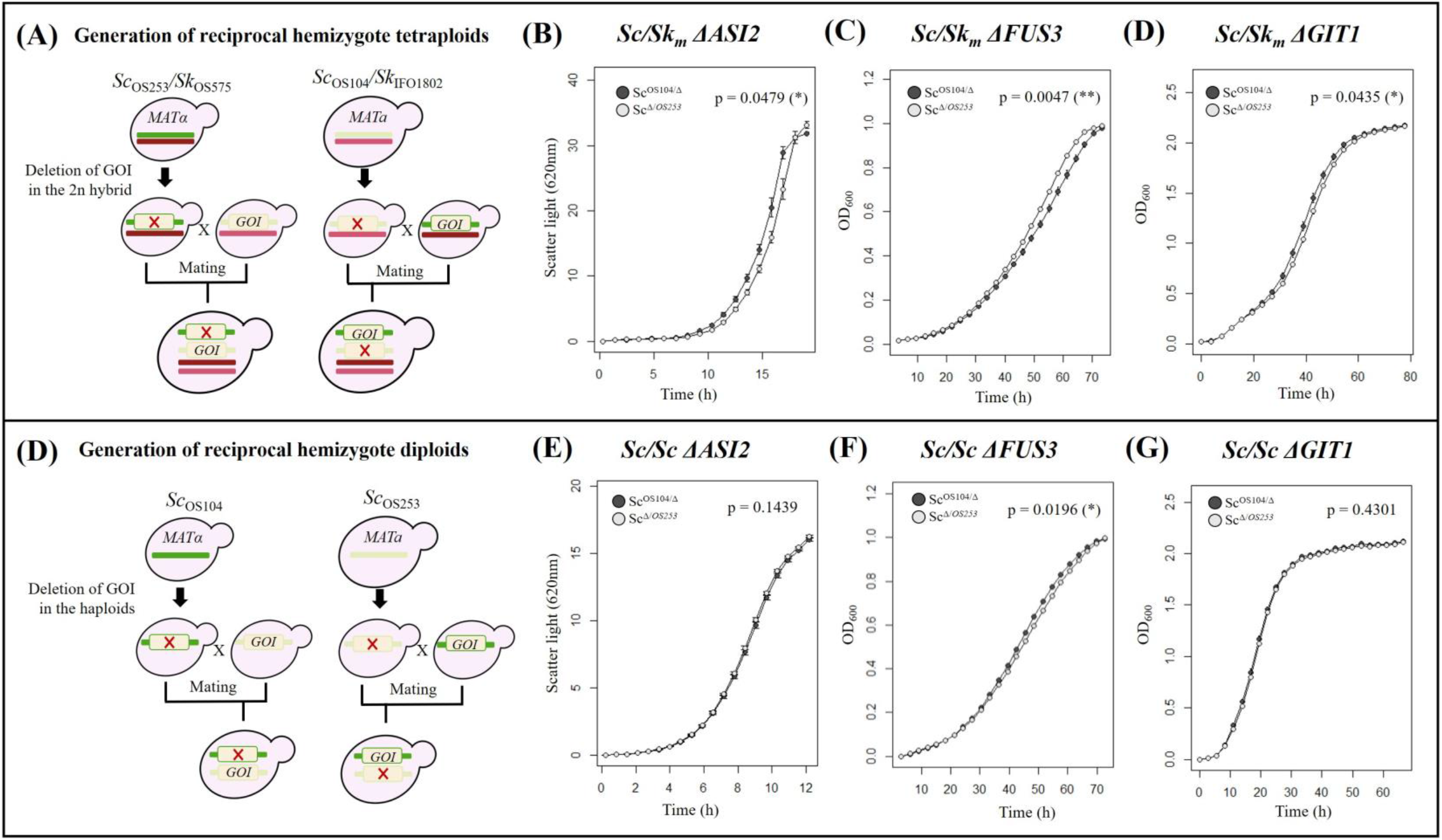
Validation of the phenotypic effect of candidate genes in inter- and intra-species hybrid background. Diagram of the construction of the reciprocal hemizygote ScSk_m_ tetraploid strains (Panel A). The gene of interest (GOI) was first deleted from the respective ScSk diploid hybrids. The engineered 2n hybrids were crossed to construct the reciprocal hemizygote tetraploid strains. Growth curves of ScSk_m_ reciprocal hemizygotes for the *ASI2*, *FUS3* and *GIT1* genes are shown in Panels B, C and D, respectively. Diagram of the construction of the reciprocal hemizygote diploid strains (Panel D). The gene of interest (GOI) was deleted from Sc_OS104_ and Sc_OS253_. The engineered haploid strains were crossed to construct the hemizygote diploids. Growth curves of Sc OS104/OS253 reciprocal hemizygotes for the *ASI2*, *FUS3* and *GIT1* are shown in Panels E, F and G respectively. Fitness assays were performed in YPD supplemented with 0.3% acetic acid (B, E), YPD at 10°C (C, F) and YP + maltose 15% (D, G) as outlined in Material and Methods. Significance difference between the integral area of the reciprocal hemizygotes is shown as p-value assessed by t-test.

### Inter-species hybrids generate new QTLs not present in the parent species

Finally, we investigated whether the phenotypic effect of the *ASI2*, *FUS3* and *GIT1*, allelic variants was exclusive to the inter-species hybrid background rather than to the intra-species strain variation (Figure 5, Panel E, F, G, H). In this case, the validation was performed through RHA on *S. cerevisiae* Sc^OS104^/Sc^*OS253*^ diploids to allow us to tease apart hybrid-dependent QTLs in inter-species crosses. No significant difference in fitness between the allelic variants was observed for both *ASI2* and *GIT1* alleles, indicating that such QTLs are exclusive to inter-species hybrid background (Table S14). In the OS104/OS253 diploids, *FUS3* alleles showed a significant difference in the integral area of the curve (p value = 0.0196; Table S14) and growth rate (p value = 0.0014, Table S14). This phenotypic variation is however opposite to that one observed in Sc/Sk_m_ tetraploids, suggesting that also in this case the allelic differences are influenced by the inter-species hybrid genomes. Moreover, a temperature-QTL including *FUS3* was also mapped in Sc/Sj_m_ hybrids suggesting that the phenotypic effect may be indeed hybrid background independent.

## Conclusions

*Saccharomyces* yeast hybrids have now entered the realm of classical genetic analysis as well as molecular genetic analysis. Previous studies have created variants of many inter-species hybrids and some even have been able to complete one round of meiotic recombination allowing some linkage analysis of genetic variants associated with traits of interest. Here we take this one step further by overcoming hybrid sterility in ways that allow continuous sequential crossing resulting in advanced intercross lines that bring in the full power of breeding genetics and quantitative genetic analysis of hybrid traits. We demonstrate that multiple traits of interest can be analysed with the same sensitivity and resolution as performed for intra-species studies. Moreover, we compare different QTL analysis approaches and can advise that the Multipool approach is more efficient for detecting most QTL than pools of highly selected subpopulations. The multipool approach will not resolve tightly linked sets of QTN within a QTL, such as the *ASI2* alleles likely to be linked to other alleles of opposite phenotypic effect. In such cases the highly selected pool can identify these complex situations by enriching for rare recombinant haplotypes in the region (Liti and Louis 2012). With the sterility of hybrids overcome, we have shown that the hybrid situation is even more complex than the complex trait analysis within a species. First, as has been seen previously, the species-specific mitochondria inherited in a hybrid has profound effects on phenotypic variation (Albertin et al, 2013; Hewitt et al, 2020) with various mitochondrial-nuclear interactions possible (Lee et al., 2008). Second and even more profound is that new QTL are generated in hybrids – that is allelic variation that has no phenotypic consequences in a parent species, has phenotypic consequences in the hybrid. The potential, therefore, for exploiting natural genetic variation in developing new hybrids, is greater than expected and bodes well for future advances in yeast breeding for improvement.

## Materials and Methods

### Strains, growth conditions and sporulation

The complete list of diploid strains used in this study is shown in Table S15. These strains were chosen for the creation of *de novo* inter-species and intra-species hybrids. Yeast strains were routinely cultured in YPD medium (1% yeast extract, 2% peptone, and 2% glucose, Formedium, Norfolk, UK). To select for the drug resistance markers, YPD medium was supplemented either with 300 μg/ml geneticin, 300 μg/ml hygromycin B, 100 μg/ml nourseothricin or 10 μg/mL bleomycin.

### Construction of genetically stable haploid strains

Genetically stable haploid *S. cerevisiae* strains used to make the hybrids were obtained from the Louis lab (Cubillos et al. 2009) and from derivatives of these (Louvel et al. 2014). *S. jurei* haploid strain was constructed previously in Delneri lab (Naseeb et al. 2017). *S. uvarum*, *S. eubayanus* and *S. kudriavzevii* haploid strains were engineered in this study. All the haploid strains used in this study are listed in Table S16. Prototrophic diploid strains were made heterothallic by knocking out the *HO* gene using standard PCR-mediated gene deletion strategy containing drug resistance cassettes (Guldener et al. 1996). The marker cassettes were amplified from plasmids listed in Table S17. Plasmids pAG32, pZC1, pZC2 and pZC4 were used to create stable haploids, pS30 pFA6-KanMX4 was used to create diploid maters in *S. eubayanus* and *S. uvarum* strains, whilst pFP18 was used to this effect in *S. cerevisiae* strains. A standard PEG/LiAc heat-shock protocol was used for transformation (Gietz and Schiestl 2007). The *HO* gene deletion was verified by diagnostic colony PCR using gene specific and cassette-specific primers. The primers used for amplification of gene knock-out cassettes and for verification of gene deletions are listed in Table S18. The heterozygote *HO/ho*Δ diploid strains were sporulated and tetrads were dissected to obtain stable *Mat***a** and *Matα ho*Δ haploid strains.

### Generating Tetraploid Hybrids and Sporulation

Mass mating, sporulation and tetrad dissection were conducted by following standard protocols (Sherman and Hicks 1991). The sporulation of the strains was carried out in sporulation medium (potassium acetate 1%, agar 2%) for five days at 21°C. Two approaches to generating tetraploids of hybrids were used (see Figure 1). One, developed by Greig et al. (2002) started with diploid hybrids between *S. cerevisiae* and other *Saccharomyces* species (inter-species diploids). In these the *S. cerevisiae* copy of the *MAT* locus was deleted creating a diploid that behaved as a haploid. In the study of Greig et al. (2002) these were *HO*, so they switched and mated creating tetraploids with the two remaining MAT loci coming from the non-cerevisiae parent. Here we used *ho*Δ haploids to start with and created two versions of diploid hybrids where in one the *S. cerevisiae* parent had one mating type while the other had the opposite. In addition, two different *S. cerevisiae* strains and two of the other parent species were used to incorporate genetic diversity in each tetraploid. The *S. cerevisiae MAT* locus in each was deleted in the same way as in Greig et al. (2002) and the resultant mating diploid hybrids were mated generating tetraploids, heterozygous for many SNPs across both genomes. In this method every meiotic spore will be a diploid hybrid that can mate, allowing further rounds of mating and meiosis. The second method started with heterozygous diploids of each species (intra-species diploids). Here one of the two *MAT* loci is deleted and species diploids of opposite mating type are mated generating tetraploids, also heterozygous for many SNPs across both genomes. In this case, meiotic spores will all be diploid hybrids but will be a mixture of non-maters and maters, still allowing for continuous rounds of mating and meiosis. In generating the various combinations of diploids to start with, the *MAT* locus was analyzed by the PCR method described previously (Huxley et al. 1990). The species of diploid spores was determined by species specific PCR (Muir et al. 2011). The primers used are listed in supplementary Table S18. The cycling condition of the PCR reaction to amplify both regions are as follows: 3 min at 95°C for the first cycle, then 1 min at 95°C, 1 min at 55°C, 3 min at 72°C for 35 cycle, and 5 min at 72°C for the last cycle.

### Multigenerational advanced intercross lines

Tetraploids from either method one or two (above), were subjected to sporulation conditions as described. After 5-7 days of sporulation, tetrads were isolated for dissection (as above) for assessing spore viability and phenotypic variation or random spore generation for further rounds of mating. For random spores, asci were washed from the sporulation plate with the help of glass beads and water. Suspended asci were pipetted into Eppendorf tubes and washed once with water and resuspended in 0.5ml water. 0.5ml of di-ethyl ether was added and the mixture vortexed for 10 minutes to kill any vegetative, unsporulated cells (Dawes and Hardie 1974). These were then microfuged and the supernatant removed. The spore pellets were washed twice in water and resuspended in dissection buffer and treated with Zymolyase 20T (1/10 volume of 10mg/ml solution) at 37°C for 30 minutes before washing and plating onto YPD media to allow germination and mating. After growth for 24 hours, the lawn of cells is replica plated onto new sporulation media and the cycle starts again. In method one, all cells that can sporulate in subsequent rounds are tetraploids generating diploid hybrid progeny. In method two, approximately 1/3 of the sporulating cells will be diploid non-mating hybrids from the previous round and their spores will be non-viable and not enter the next round. The rest of the sporulating cells will be tetraploids generating the same mixture of mater and non-mater diploid hybrid progeny.

### Analysis of mitochondrial origin in hybrids

For each diploid mater, a petite version was generated by exposure to ethidium bromide (EtBr) (Goldring et al. 1971). Isolates were seeded at an approximate density of 300 individuals per plate. A 3μl drop of EtBr (10 mg/ml) was spotted onto the centre of each plate. A ring around the spot formed where all cells were killed due to the toxic effects of EtBr. Surrounding this kill zone, a ring of petite colonies form. Loss of mitochondria enables petites to grow faster than colonies with functional mitochondria, in the presence of EtBr. These individuals were confirmed as petites by their inability to grow on YEPEG plates containing ethanol and glycerol, non-fermentable carbon sources (Day 2013). The specific mitotype was identified by amplifying the *COX2* and *COX3* genes through colony PCR as described previously (Belloch et al. 2000 and Hewitt et al 2020). The primers used for the amplification of these genes are listed in Table S18.

### Phenotypic Assays

A high-throughput spot assay was performed using Singer ROTOR HDA robot (Singer Instruments, UK) as mentioned previously (Naseeb et al. 2018). The fitness of ~384 hybrid spores was assessed at five different temperatures *i.e.* 10°C, 16°C, 23°C, 30°C and 37°C, under different carbon sources at 30°C *i.e.* YPA + 10% & 15% maltose, YPA + 10% & 15% fructose, YPA + 10% & 15% sucrose, YPA + 10% & 15% galactose, YPA + 30% & 35% glucose and under different environmental stressors at 30°C *i.e.* YPAD + 6% & 10% ethanol, YPAD + 0.3% & 0.5% acetic acid, YPAD + 4 mM hydrogen peroxide, YPAD + 50 mM levulinic acid and YPA + 10% & 15% glycerol.

Fitness analysis was done following two different strategies. In strategy one, high-resolution images of phenotypic plates were taken using Phenobooth after 3 days of incubation (Singer Instruments, UK). The colony sizes were calculated in pixels using Phenosuite software (Singer Instruments, UK) and the heat maps of the phenotypic behaviours were constructed using the R shiny app (https://kobchai-180shinyapps01.shinyapps.io/heatmap_construction/). In strategy two, phenotyping was performed using the PHENOS platform (Barton et al. 2018). Change in absorbance (600 nm) was measured using the FLUOstar Omega plate reader (BMG Labtech). Prior to seeding cells on selection plates, a blank reading was taken, with values subtracted from each time course reading. Plates were incubated at 30°C (except for temperature selection plates) and absorbance was measured every 20 minutes over the course of three days. Individuals were ranked by the maximum change in absorbance, after normalising to the maximum change in absorbance under control conditions.

### Assessing genetic variation using Illumina sequencing

The engineered hybrid progeny were sequenced to determine the genetic variation responsible for particular traits of interest. Each parental genome in the hybrid was sequenced previously (Naseeb et al. 2018) or in this study, enabling us to determine the inheritance of every segment of the resulting diploid hybrid progeny genomes. The sequencing was done to 100-120X coverage on the Illumina platform by pooling individual diploid hybrid spores with appropriate phenotypes and sequencing the pool for comparison to the original population. Two different pooling strategies were followed. In strategy one, from each selective media plate the top twenty performing F12 individuals, with the highest fitness, and the twenty lowest performing F12 individuals were picked for pooling. In second strategy, a pool of 1 * 10^8^ F12 cells were seeded onto each selection condition as well as the YPD control. Selection conditions were prepared so that only the top 0.1 – 1 % of the pool would be capable of growth. After four days the cells were washed off the selection plates and collected in H_2_O. 50 % of the cell population was stored in glycerol at −80C, with DNA extracted for sequencing of the other 50 %. QTL regions associated with the phenotype were identified by analyzing the changes in frequencies of SNP alleles across the genomes.

### Sequencing, Mapping and Variant Calling

The hybrids were sequenced by the Earlham Institute and GTCF in the University of Manchester. Paired-end raw Illumina sequence reads in fastq format were quality checked through FastQC 0.11.5 (Andrews 2010) and trimmed through Trimmomatic (Bolger, et al., 2014). Filtered reads were aligned to a combined reference genome with parental species (*S. cerevisiae* YP128 (Yue et al. 2017) concatenated with *S. eubayanus* CBS12357 (Libkind et al. 2011) / *S. jurei* D5088 (Naseeb et al. 2018) / *S. kudriavzevii* (assembled genome) using bwa/0.7.16a (H. Li and Durbin 2009). The SPAdes assembler 3.9.0 was applied on the filtered reads of *S. kudriavzevii* IFO 1802 after quality checking and trimming for assembly. Assembly quality was assessed through QUAST/4.3. Local realignment was then performed to minimise the number of mismatching bases through RealignerTargetCreator and IndelRealigner through GATK 3.8. The MarkDuplicates tool was then used with picard/2.6.0 for removing the optical duplicates to control the alignment quality for variant analysis. Samtools was applied on step with raw bam files for sorting and indexing (H. Li et al. 2009). Variant calling was then applied on the aligned reads using freebayes/1.0.2 (Garrison E 2012) with ploidy setting at 1 with the combined reference, --min-mapping-quality 30 --min-base-quality 20 --no- mnps. Variant calling outputs in vcd files were then filtered with only SNP results and transferred to csv files that included information of CHROM, POS, REF, ALT, AO, RO where CHROM is the chromosome where each SNP is located, POS is the physical position in reference genome, REF is the base of reference genome ALT is the base of variants, AO is the allele depth of alternatives and RO is the allele depth that have same base with reference. Pool SNPs were further compared to the SNPs among the founders through an R script. Reads depths below 10 were excluded. For each pool, matching allele files were stored separately by chromosome in txt for next stage allele frequency tests between high fitness and low fitness. The Variant calling pipeline is illustrated in Figure S4. The analysis for mapping and variant calling were performed on HPC service with qsub files. SNP filtering to genotype was analysed in R with scripts.

### Gene ontology and SIFT analysis

Gene ontology (GO) terms were determined using the GO Term Finder tool of Saccharomyces Genome Database (SGD) with the Bonferroni correction for multiple hypothesis and a p-value cutoff of 0.01.

Potential causal genes were analysed with the Sorting Intolerant from Tolerant (SIFT) algorithm to assess if amino-acids variants were predicted to influence the protein function. SIFT analysis were conducted using data from Bergstrom et al. 2014 on the *S. cerevisiae* strains OS3, OS104 and OS253, and the tool mutfunc (Wagih et al. 2018).

### Validation of candidate genes through reciprocal hemizygosity analysis

The candidate genes selected for reciprocal-hemizygosity analysis were chosen based on their LOD score and on gene ontology studies carried out with YeastMine (Balakrishnan et al. 2012) and Funspec (Robinson et al. 2002) to prioritize terms connected to the phenotypic trait tested.

To dissect the mapped QTL regions and confirm the effect of candidate genes in determining the phenotypic fitness, we performed reciprocal-hemizygosity analysis (Steinmetz et al. 2002) on selected candidate genes: *ASI2*, *FUS3* and *GIT1*. PCR mediated deletion of the target genes was performed in the engineered *S. cerevisiae/S. kudriavzevii* hybrid (OS253/OS575 and OS104/IF01802), to delete each *S. cerevisiae* allele. The transformants were then mass mated together to construct the reciprocal hemizygotes hybrid and selected in triple drug plates (300 μg/ml geneticin, 100 μg/ml nourseothricin or 10 μg/mL bleomycin) (Table S18).

To investigate if the effect of the allelic variants was exclusive to the inter-species hybrid background rather than to the intra-species strain, PCR-mediated deletion was used to generate null mutants of the parental *S. cerevisiae* haploid strains (OS253 and OS104) for the candidate genes. The transformed haploid strains were them mass mated together to construct the reciprocal hemizygotes hybrids and selected in SD +ura +leu with 100 μg/ml nourseothric or 300 μg/ml geneticin (Table S19). Successful gene deletions were confirmed by PCR. All the primers used for the construction of the strains are listed in Table S18.

The fitness of the *S. cerevisiae* allelic variants of *FUS3*, *GIT1* and *ASI2* was assessed both in the hemizygous *S. cerevisiae / S. kudriavzevii* tetraploids and in the hemizygous *S. cerevisiae* diploids. The *FUS3* and *GIT1* allelic variants was assessed with a BMG FLUOstar OPTIMA Microplate 466 Reader as previously described (Naseeb and Delneri, 2012) in YPD at 10°C, and at 25°C in YPD + 15% maltose, respectively. The fitness of *ASI2* hemizygous hybrids was assessed with the BioLector (m2p Labs, GmbH). A 48-well flowerplate with transparent glass bottom was inoculated with overnight cultures at a starting OD_600_ of 0.1. Every culture was run in triplicates in 1.5 ml of YPD + 0.3% acetic acid. The flowerplate was incubated at 30°C in the BioLector with orbital shaking of 800 rpm and oxygenation was maintained at 20%. Scatter light readings, to measure the cell density, were taken from the bottom of each well every 8 min with a gain of 15 and 25. The growth characteristics of the plate reader experiments were assessed with the R package Growthcurver using K as maximum biomass, *r* as maximum growth rate, *auc_l* as integral area (Sprouffske and Wagner 2016) and *Tmid* as the time at which the population density reaches *1/K*.

**Table 1.** Number of QTL regions detected in *S. cerevisiae/S. jurei*, *S. cerevisiae/S. kudriavzevii* and *S. cerevisiae/S. eubayanus* F12 segregants via Multipool strategy

|  | Number of QTL regions detected |  |  |  | Number of genes within the QTL regions |  |  |  |
| --- | --- | --- | --- | --- | --- | --- | --- | --- |
|  | Sc<sub>m</sub>Sj Hybrid |  | ScSj<sub>m</sub> Hybrid |  | Sc<sub>m</sub>Sj Hybrid |  | ScSj<sub>m</sub> Hybrid |  |
|  | Sc genome | Sj genome | Sc genome | Sj genome | Sc genome | Sj genome | Sc genome | Sj genome |
| Low temperature (16°C) | 4 | 13 | 17 | 20 | 59 | 269 | 140 | 264 |
| Maltose | 8 | 7 | 12 | 9 | 89 | 80 | 111 | 131 |
| Acetic acid | 7 | 8 | 8 | 6 | 74 | 159 | 87 | 77 |
|  | Sc<sub>m</sub>Sk Hybrid |  | ScSk<sub>m</sub> Hybrid |  | Sc<sub>m</sub>Sk Hybrid |  | ScSk<sub>m</sub> Hybrid |  |
|  | Sc genome | Sk genome | Sc genome | Sk genome | Sc genome | Sk genome | Sc genome | Sk genome |
| Low temperature (16°C) | 33 | 11 | 20 | 25 | 243 | 64 | 171 | 95 |
| Maltose | 38 | 32 | 24 | 24 | 243 | 192 | 233 | 120 |
| Acetic acid | 24 | 28 | 19 | 10 | 164 | 200 | 188 | 39 |
|  | Sc<sub>m</sub>Se Hybrid |  | ScSe<sub>m</sub> Hybrid |  | Sc<sub>m</sub>Se Hybrid |  | ScSe<sub>m</sub> Hybrid |  |
|  | Sc genome | Se genome | Sc genome | Se genome | Sc genome | Se genome | Sc genome | Se genome |
| Very low temperature (4°C) | 13 | 8 | 13 | 11 | 188 | 83 | 212 | 121 |
| High temperature (40°C) | 18 | 10 | 26 | 13 | 82 | 76 | 137 | 173 |
| Ethanol | 15 | 8 | 26 | 14 | 137 | 120 | 232 | 173 |

## Acknowledgments

The authors wish to thank Gianni Liti, Chris Powell, Philippe Malcorps and Stewart Wilkinson for useful discussions, and Ian Donaldson for initial bioinformatic support. Sequencing was done through Genomic Services at the Earlham Institute and Genomic Technologies Core Facility at the University of Manchester. This work was funded by a BBSRC grant to EJL (BB/L022508/1) and DD (BB/L021471/1) in collaboration with SAB-Miller and AB-InBev; FV is supported by H2020-MSCA-ITN-2017 grant to DD (764364; https://cordis.europa.eu/project/id/764364). AR has been supported by BBSRC-CASE studentship to EJL (BB/L017229/1).

## Supplementary Table Legends

**Table S1.** List of inter-species diploid hybrids constructed.

**Table S2.** List of intra-species diploid hybrids constructed.

**Table S3.** List of tetraploid hybrids constructed.

**Table S4.** Spore viability of engineered tetraploid and diploid hybrids.

**Table S5.** Quartile coefficient of dispersion of the F1 and F12 segregants colony size by condition.

**Table S6.** Total number of QTLs identified in the F12 segregants in all conditions using the Multipool strategy.

**Table S7.** List of potential causal genes identified in hybrids analysed via Multipool.

**Table S8.** List of potential causal genes identified in the *S. cerevisiae* genome of hybrids analysed via Multipool with non-synonymous SNPs predicted to be tolerated or deleterious by SIFT analysis.

**Table S9**. List of pleiotropic QTLs in hybrid diploid progeny.

**Table S10**. Total number of QTLs identified in the F12 segregants in all conditions using the Pooled selection strategy.

**Table S11.** List of potential causal genes identified in hybrids analysed via Pooled selection.

**Table S12**. List of overlapping QTLs intervals identified in different hybrids or species.

**Table S13.** List of QTL regions identified in *S. cerevisiae*/*S. kudriavzevii* F12 selected for RHA.

**Table S14:** Growth parameters of reciprocal hemizygotes for the *ASI2*, *FUS3* and *GIT1* gene in inter- and intra- species hybrid background.

**Table S15.** List of strains used in this study and their origin.

**Table S16.** List of haploid strains and their genotypes.

**Table S17.** List of plasmids used for strain construction.

**Table S18.** The set of primers used for gene deletion, species identification and mitochondrial gene amplification.

**Table S19**: Strains used for reciprocal hemizygosity analysis.

## Supplementary Figure Legends

**Figure S1**. **Scatter plots of fitness of F1 diploid progeny for***S. cerevisiae/S. jurei* **and***S. cerevisiae/S. kudriavzevii* **hybrids. A)** The dot plots represent the colony sizes of *S. cerevisiae/S. jurei* (A) and *S. cerevisiae × S. kudriavzevii* (B) hybrids at different temperatures, carbon sources and stressors. Each dot represents the fitness of a F2 progeny. Different mitotypes are represented in black for *S. cerevisiae*, in red for the other parental species and in green for recombinant mitotypes.

**Figure S2. Heat map representing phenotypic fitness of F1 diploid progeny for***S. cerevisiae/S. jurei* **hybrids.** Fitness was scored at five different temperatures and in response to different environmental stressors at 30°C for 4n hybrid parents Sc/Sj carrying different mitochondria and their diploid progeny. Phenotypes are represented with colony sizes calculated as pixels and coloured according to the scale, with light yellow and dark blue colours representing the lowest and highest growth respectively.

**Figure S3**. **Scatter plots of fitness of F12 diploid progeny for***S. cerevisiae/S. jurei, S. cerevisiae/S. kudriavzevii* **and***S. cerevisiae/S. eubayanus* **hybrids. A)** The dot plots represent the colony sizes of *S. cerevisiae/S. jurei* (A and B), *S. cerevisiae × S. kudriavzevii* (C and D) and *S. cerevisiae/S. eubayanus* (E and F) hybrids at different temperatures, carbon sources and stressors. Each dot represents the fitness of a F12 progeny.

**Figure S4**. **Variant calling pipeline**

