## Supplementary figures and images for "Restoring fertility in yeast hybrids: breeding and quantitative genetics of beneficial traits"

### Figure S1 (Sc+Sj & Sc+Sk F2 fitness).jpg

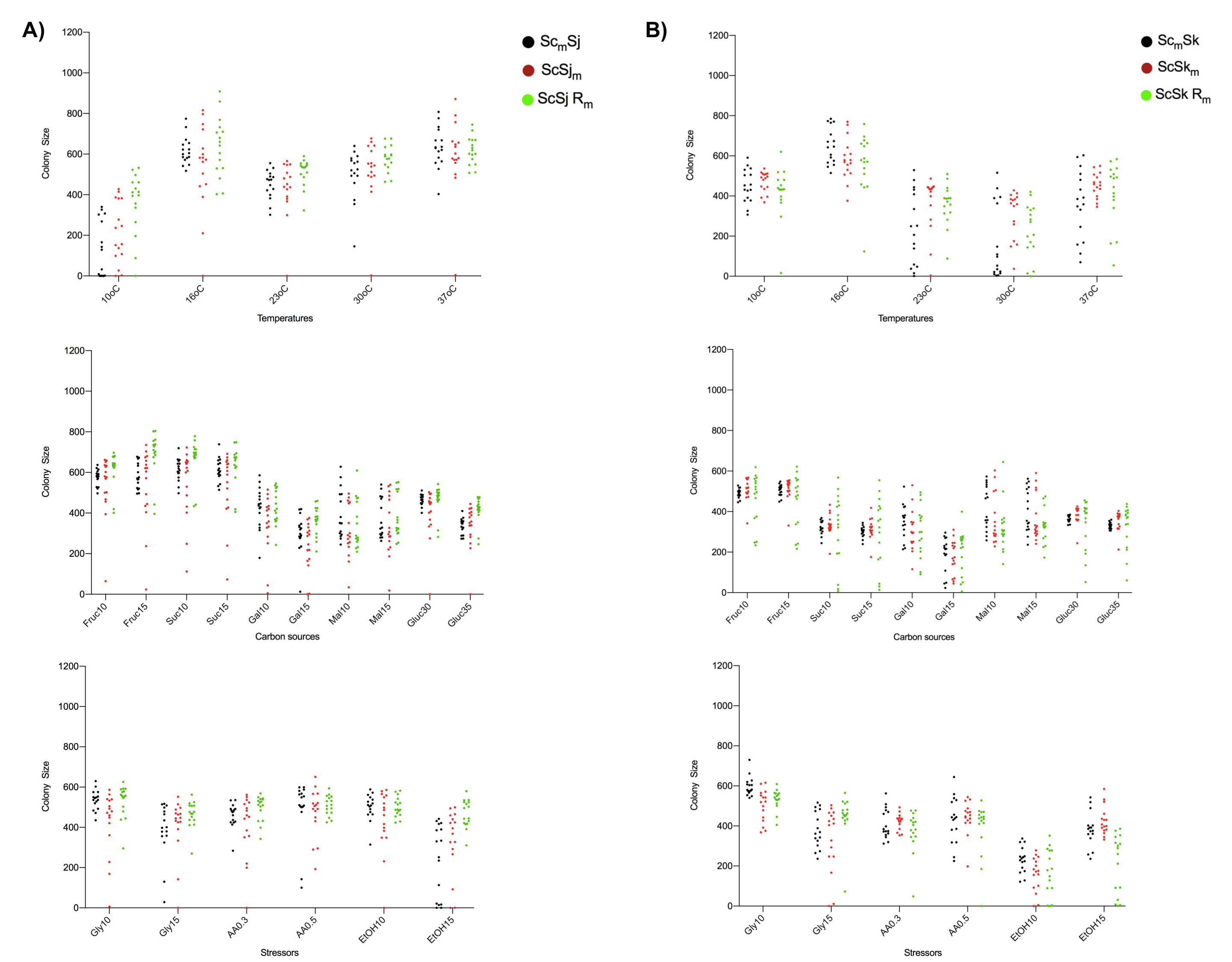

### Figure S2_F2 Sc+Sj heatmap.jpg

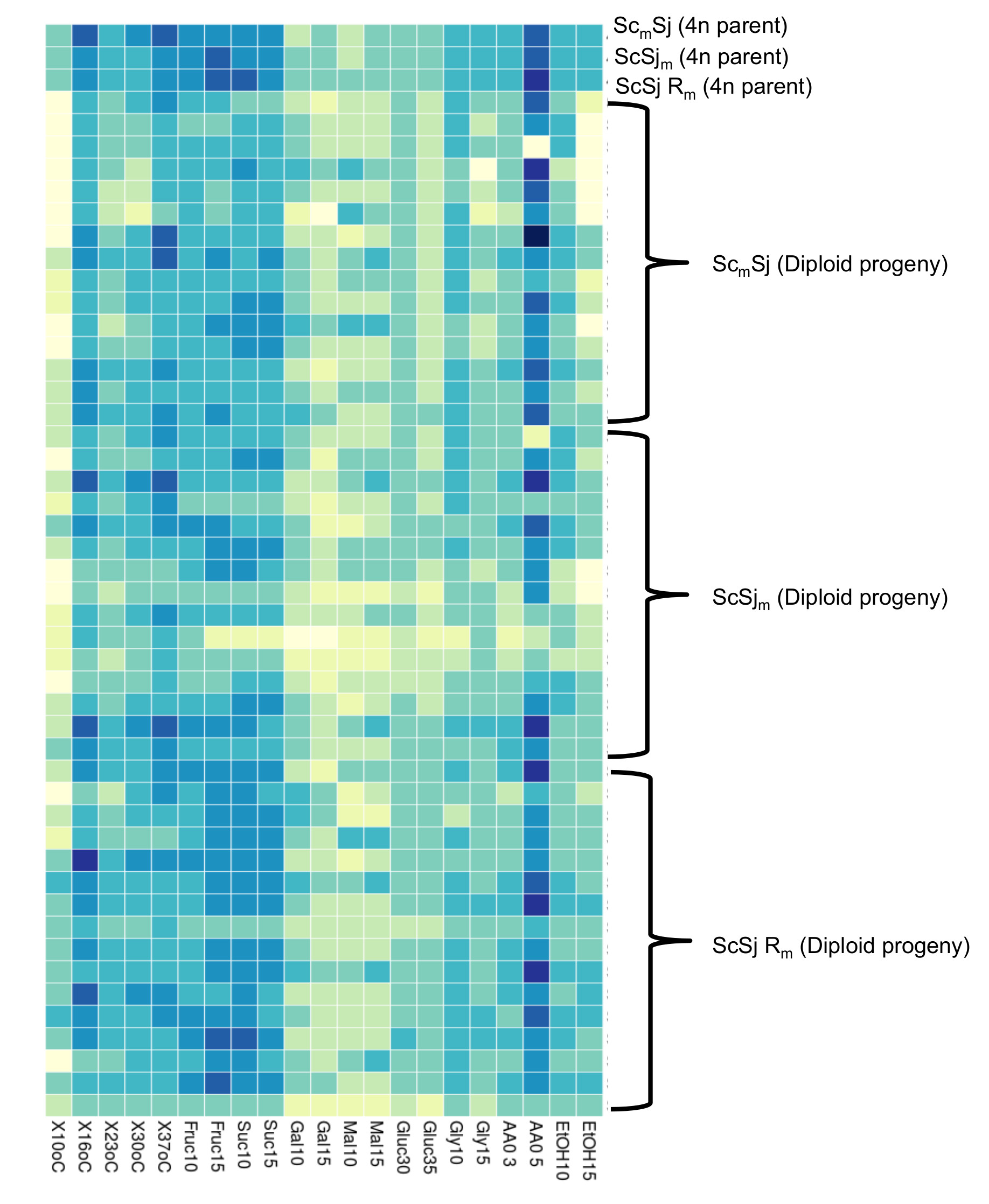

### Figure S3 (Actual colony size F12 fitness).jpg

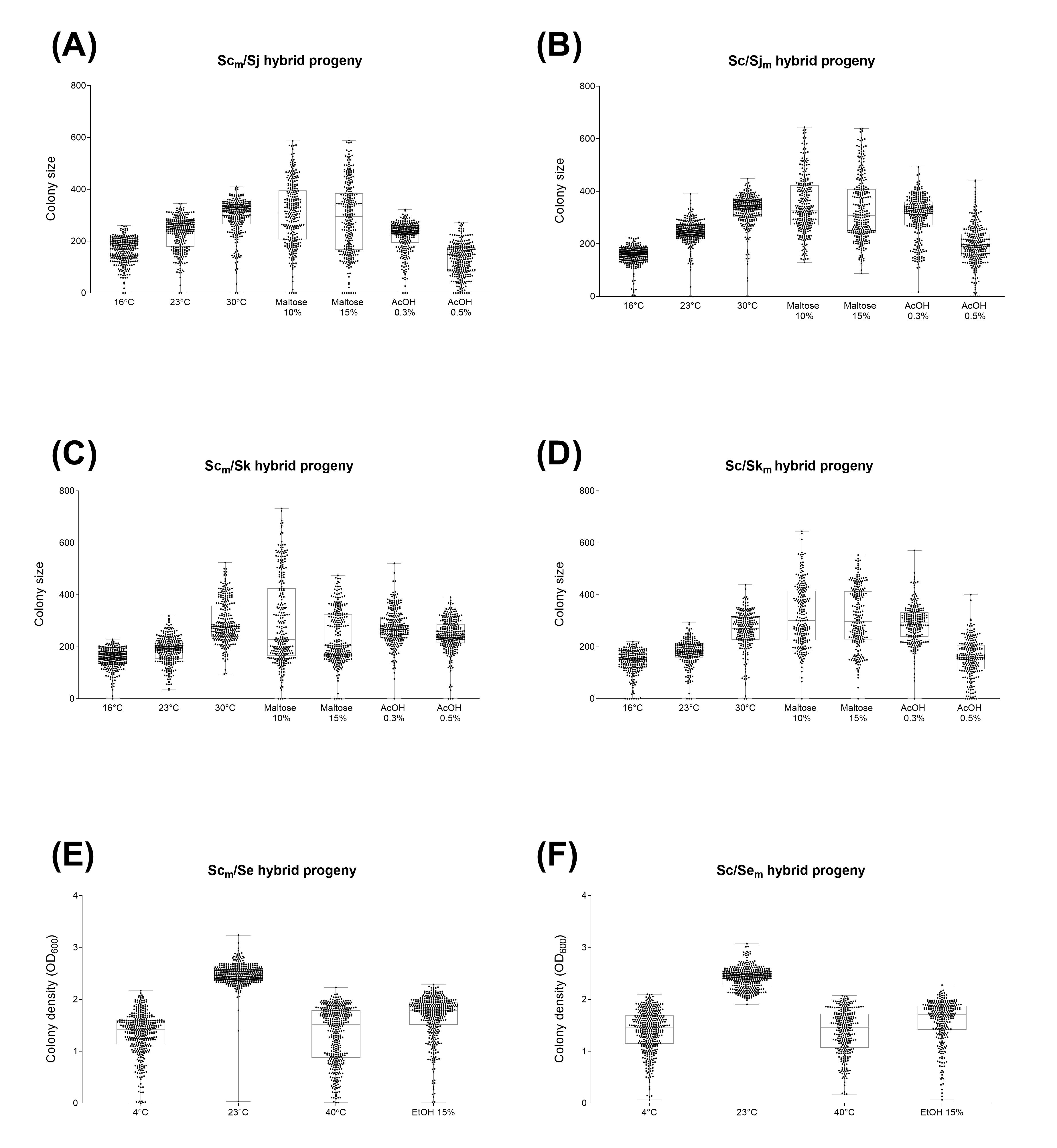

### Figure S4 - Variant calling strategy.png

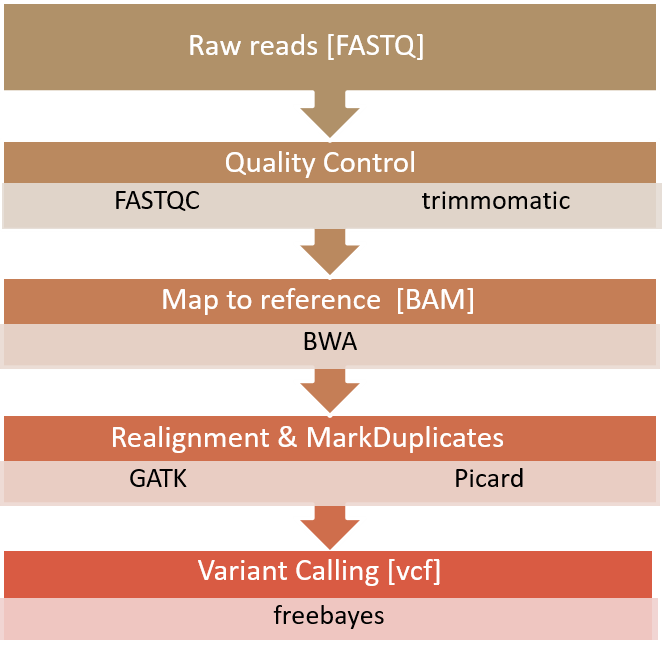
